# A Dysbiotic Gut Microbiome Suppresses Antibody Mediated-Protection Against *Vibrio cholerae*

**DOI:** 10.1101/730796

**Authors:** John Macbeth, Ansel Hsiao

**Affiliations:** Department of Microbiology and Plant Pathology, University of California, Riverside, Riverside, California, USA; Division of Biomedical Sciences, University of California, Riverside, Riverside, California, USA

**Author notes:** Address correspondence to: Ansel Hsiao.

## Abstract

*Vibrio cholerae* is the etiologic agent of cholera, a severe diarrheal disease that represents a significant burden on global health and productivity. Despite the pressing need, available preventative measures such as oral cholera vaccines exhibit highly variable protective efficacy. We hypothesized that one contributor to vaccine efficacy variability across geographical regions may be due to differences in gut microbiome, which in cholera-endemic areas is strongly and repeatedly modulated by malnutrition, cholera, and non-cholera infectious diarrhea. Here, we assemble representative model communities of either human gut microbes resembling those of healthy individuals or those of individuals recovering from diarrhea or malnutrition. We establish these communities in a murine immunization model, and show that the dysbiotic gut microbiome, commonly present in areas where malnutrition and diarrhea are common, suppresses the immune response against *Vibrio cholerae* through the action of CD4^+^ cells. Our findings suggest that the composition of the gut microbiome at time of immunization may be pivotal for providing robust immunity from oral cholera vaccines, and highlight the importance of the gut microbiome on mucosal immunization responses and vaccine development strategies.

**Importance:** Diarrhea caused by enteric bacterial pathogens is a recurring and important issue for worldwide health. Cholera, the severe watery diarrhea caused by the bacterium *Vibrio cholerae*, affects millions of people annually. Currently, there is a lack of effective preventative measures for cholera, due to the uneven performance of oral cholera vaccines. Thus, it is essential to better understand the factors that may affect vaccine efficacy. One aspect may be variations in the resident community of gut microbes, the gut microbiome, across populations living in developed versus developing regions as a function of host genetics, diet, and infection. Our findings suggest that specific structures of the gut microbiome are involved in disrupting the immune responses to *V. cholerae* vaccination.

## Introduction

*Vibrio cholerae* is the etiologic agent of cholera, a severe diarrheal disease affecting approximately 3 million people annually, resulting in over 100,000 deaths (1). *V. cholerae* preferentially colonizes the small intestine, where it releases cholera toxin (CT), which causes profuse watery diarrhea and loss of electrolytes. While the advent of oral rehydration therapy has dramatically reduced mortality from cholera, recent major outbreaks in Yemen and Haiti are reminders of the pressing global public health need to improve cholera prevention strategies. Several oral cholera vaccines (OCVs) have been developed, though they have demonstrated high variance in protective efficacy in field trials (2). While OCVs have been shown to have an efficacy of generally 80%-90% in developed countries, large field studies in developing regions range from 14-50% efficacy (3-7). Two main types of OCVs have been developed, whole-cell killed and live-attenuated. The whole-cell killed vaccine, Dukoral, consisted of inactivated strains of O1 Classical and El Tor biotype *V. cholerae* as well as recombinant cholera toxin subunit B, the non-enzymatic subunit of cholera toxin (7). Dukoral showed 50% efficacy in all age groups in a Bangladesh field trial (3), while demonstrating significant vibriocidal titers (a clinical correlate of protection) in 89% of volunteers in the US (4). Another whole-cell killed vaccine, Shanchol, is composed of various O1 Classical and El Tor strains as well as one O139 strain (7). In children aged 1-5 years in Kolkata, the vaccine had an efficacy of 43% after 5 years compared to 65% in the 5-15 years of age group (8, 9). The only OCV to be approved by the FDA in the United States, Vaxchora, comprises a live-attenuated O1 Classical Inaba strain, with 94% of the *ctxA* gene encoding the enzymatic subunit of CT deleted and replaced by a mercury-resistance cassette, while leaving the *ctxB* gene intact. While the vaccine was safe and immunogenic in US trials (10), it had a less than desirable outcome of 14% protection in a large Bangladesh field trial (6).

We hypothesized that one contributor to this high level of geographical variation in OCV efficacy may be variations in the microbial populations of the gut, the gut microbiome. Several studies have shown that gut bacterial populations can change due to diet and geography (11, 12), especially when comparing populations in the United States and Europe versus those in developing countries (13-15). While the presence of a normal murine microbiome has been implicated in antibody responses against viral vaccines (16), the effects of a human gut microbiome on responses to *V. cholerae* or other enteropathogenic bacteria have not been well determined. To begin to address this question, we have assembled human fecal bacterial communities that are representative of either bacteria that are commonly found in healthy human guts (17) or bacteria prevalent in malnutrition or diarrhea endemic regions based on existing 16S ribosomal DNA amplicon surveys (11, 18). Here, we show that dysbiotic gut bacterial populations at time of *V. cholerae* immunization dampen antigen-specific antibody responses against *V. cholerae* in a CD4+-cell-dependent manner. These findings suggest that gut bacterial composition at time of immunization may impact adaptive immune responses to *V. cholerae* and may be applicable to other enteric pathogens as well.

## Results

To begin to examine the role of human microbiome structure on vaccination outcomes, we established model human fecal communities in adult CD-1 mice. Preclinical studies in animal models are essential to understand the intricate mechanisms of vaccine protection. There are several animal models for studying *V. cholerae*, the most widely used being the infant mouse cholera model (19). However, while the suckling animals can be used to study *Vibrio* colonization and virulence, they are poorly suited for immunological studies, as the infant mouse does not have a fully developed adaptive immune system, a limitation shared by the recently developed infant rabbit model of cholera (20, 21). Others have utilized germ-free adult mice to evaluate immune responses to *V. cholerae* (22), but there may be limitations to this approach as well because there are no microbial-immune system interactions at birth, which are critical to a robust development of adaptive immunity (23). However, depletion of the murine microflora is critical to examining the effects of human microbial communities, as mouse-adapted microbes rapidly out-compete non-murine communities (24). Thus, we used antibiotics to treat adult animals (25) in order to allow the introduction of human gut communities in an immune-competent animal system. Adult CD-1 mice were treated with an antibiotic cocktail for 1 week (See Materials and Methods) and then switched to streptomycin treatment alone 3 days before the gavage (Fig. 1A). The mice were then gavaged with ∼5 × 10^9^ CFU of *V. cholerae* C6706 O1 El Tor. We observed that antibiotic treatment was critical to eliciting antibody responses against immunization (Fig. 1B). Serum vibriospecific ELISA showed that levels of IgG3 and IgM, strong complement fixing antibodies, were decreased in the NM group as compared to the DM group (Fig 1C).

**FIG 1.**
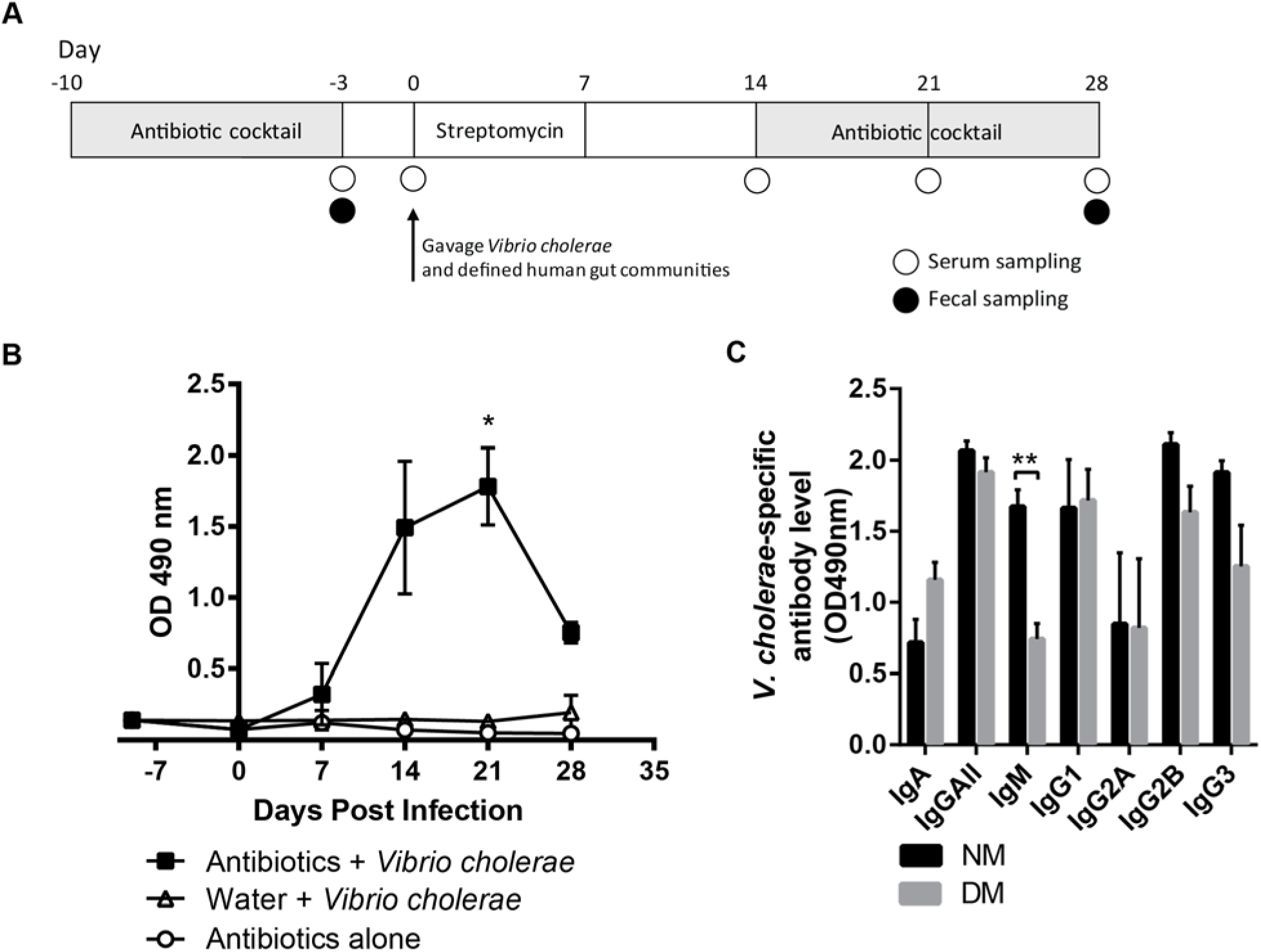
*V. cholerae-*specific antibody levels in fecal and serum samples. (A) Schematic of antibiotic treatment in SPF CD-1 mice. (B) Post-immunization vibriospecific fecal IgA increased in the antibiotic treated group as compared to other groups. (C) Serum antibody profiles against whole cell *V. cholerae* 4-weeks post-immunization. NM: normal model microbiome, DM: dysbiotic model microbiome. *, *P*<0.05, **, *P*<0.01, unpaired Student’s *t*-test.

We then moved to establish human gut communities in these antibiotic cleared mice. As a proof of principle, we constructed simplified defined communities of human gut isolates, designed on the basis of metagenomic surveys of individuals in cholera-endemic areas, but nonetheless simple enough to be easily and reproducibly deployed in numerous animal experiments. We constructed two embodiments, the NM microbiome, broadly representative of healthy microbiomes (13, 26) and the DM model microbiome, which closely resembled dysbiotic human gut microbial communities in areas affected by malnutrition and diarrhea, including cholera (11, 17, 18, 27) (Table S1).

At 2 weeks post introduction of *V. cholerae* the mice were switched to ampicillin, neomycin, and vancomycin in drinking water. We re-introduced strong antibiotic control of microbial growth in these animals for two reasons. First, we wanted to remove *V. cholerae* so that potentially varying levels of antigen present over time would not affect immune responses; germ-free adult mice show sustained colonization, while human gut microbial communities have been shown to affect colonization levels in mice (17). Shedding of *V. cholerae* in Vaxchora recipients has been minimal, and continued bacterial carriage for any vaccine is disadvantageous for safety reasons (28). In addition, we wanted to remove the introduced human microbiomes after they had had the opportunity to influence responses to *V. cholerae*, and before any potential long-term fluctuations in the community can induce immunomodulatory effects on the host. We reasoned that this approach also more closely approaches practice in vaccine deployment in humans in endemic areas. Individuals suffering from malnutrition and various forms of infectious diarrheas in Bangladesh have demonstrated a strong but transient shift in fecal microbial composition to the DM community structure (18). This temporary dysbiosis is likely to affect a substantial portion of human vaccine recipients, and has not served as an exclusion criteria for prior vaccine trials.

At 4 weeks post introduction of *V. cholerae*, serum and fecal samples were collected from immunized mice containing NM and DM human microbiomes. To evaluate the efficacy of anti-*V. cholerae* antibody responses, we used a serum vibriocidal assay, which is considered to be the best clinical correlate of protection for cholera (29-32). The vibriocidal titer is the reciprocal of the highest dilution of serum at which killing of *V. cholerae* is observed with the addition of exogenous complement. Briefly, serum from immunized animals is heat-inactivated and diluted in two-fold steps, after which a mixture of guinea pig complement and *V. cholerae* are added to each serum sample. After incubation, *V. cholerae* growth is observed by plating on selective solid medium. In humans, a clinically successful seroconversion as a result of vaccination is defined as a more than four-fold rise in serum vibriocidal titers compared to baseline pre-immune titer over two weeks, although there is no defined titer at which protection can be considered to be definitively achieved (33). We observed that serum from animals bearing the (DM) microbiome at time of infection exhibited a statistically significant reduction in serum vibriocidal activity compared to that from animals immunized in the presence of the (NM) microbiome (Fig 2), suggesting that the presence of members of the dysbiotic community at time of immunization may hinder the development of a robust serum antibody response.

**FIG 2.**
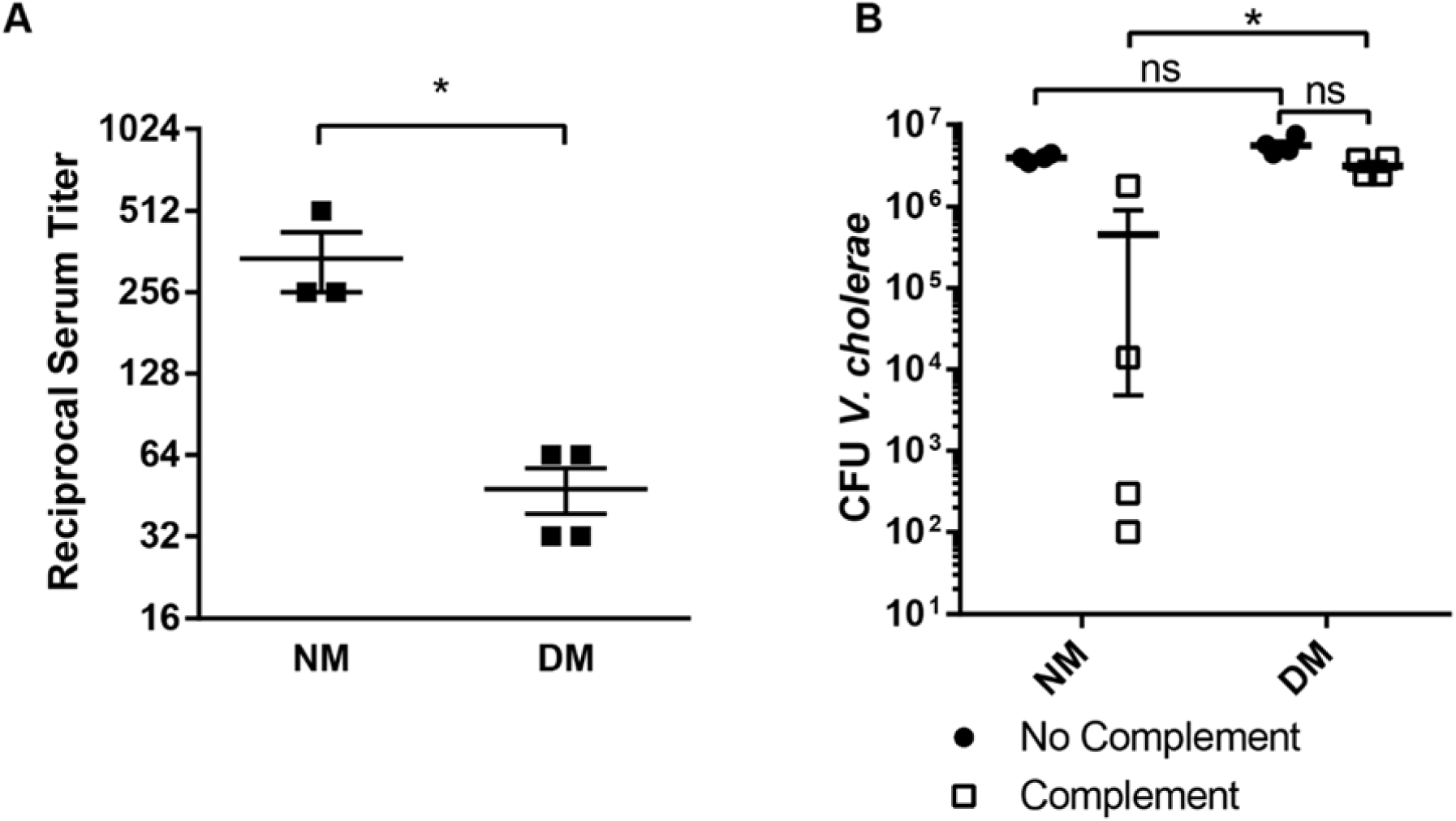
Serum vibriocidal activity after immunization in presence of normal (NM) and dysbiotic (DM) model human microbiomes in antibiotic-cleared CD-1 mice. (A) Serum vibriocidal titer 4-weeks post-immunization. (B) Levels of *V. cholerae* survival after incubation with serum antibody and exogenous complement. *, *P*<0.05, Mann-Whitney *U* test.

Although the vibriocidal assay represents a good correlate of protection in humans, we used a passive protection assay to determine a more functional measure of the activity of antibody generated by immunized animals. While serum vibriocidal activity is mostly due to the action of IgG and IgM, protection from infection is thought to be primarily mediated by secreted antibodies, especially secretory antibodies at the site of infection, i.e. the intestinal mucosa. During the course of infection, class-switching to IgA and the secretion of antigen-specific secretory IgA (s-IgA) serves as the main means of protection by binding to *V. cholerae* and preventing pathogen access to epithelium, and neutralizing cholera toxin (34). A recent study highlights the capacity of a monoclonal IgA antibody to inhibit *V. cholerae* motility, preventing access to the intestinal epithelium (35). IgA is not likely to be well reflected in serum; the bulk of IgA in the body is secretory IgA (s-IgA) secreted in gram quantities onto the mucosa (36). To obtain this secreted antibody from vaccinated animals, fecal immunoglobulin from both NM and DM animals was purified using a Protein L Purification Kit (Pierce). These antibody pools were predominantly IgA with low levels of IgM (Fig 3A). Purified Ig from both groups was combined with *V. cholerae* grown overnight and incubated for 1 hour before being gavaged into 4-day old infant CD-1 mice. Suckling animals were used as conventionally-reared adult animals are highly resistant to *V. cholerae* colonization (37, 38). After 18 hours of infection, the small intestines were homogenized and plated on selective medium. We observed that pre-treatment with antibody from animals bearing the dysbiotic microbiome led to colonization nearly 2-log greater than pre-infection treatment with antibody from animals with the NM microbiome (Fig. 3B). Taken together, these findings suggested that oral immunization in a DM microbiome context lead to a significantly less effective anti-*Vibrio* antibody response.

**FIG 3.**
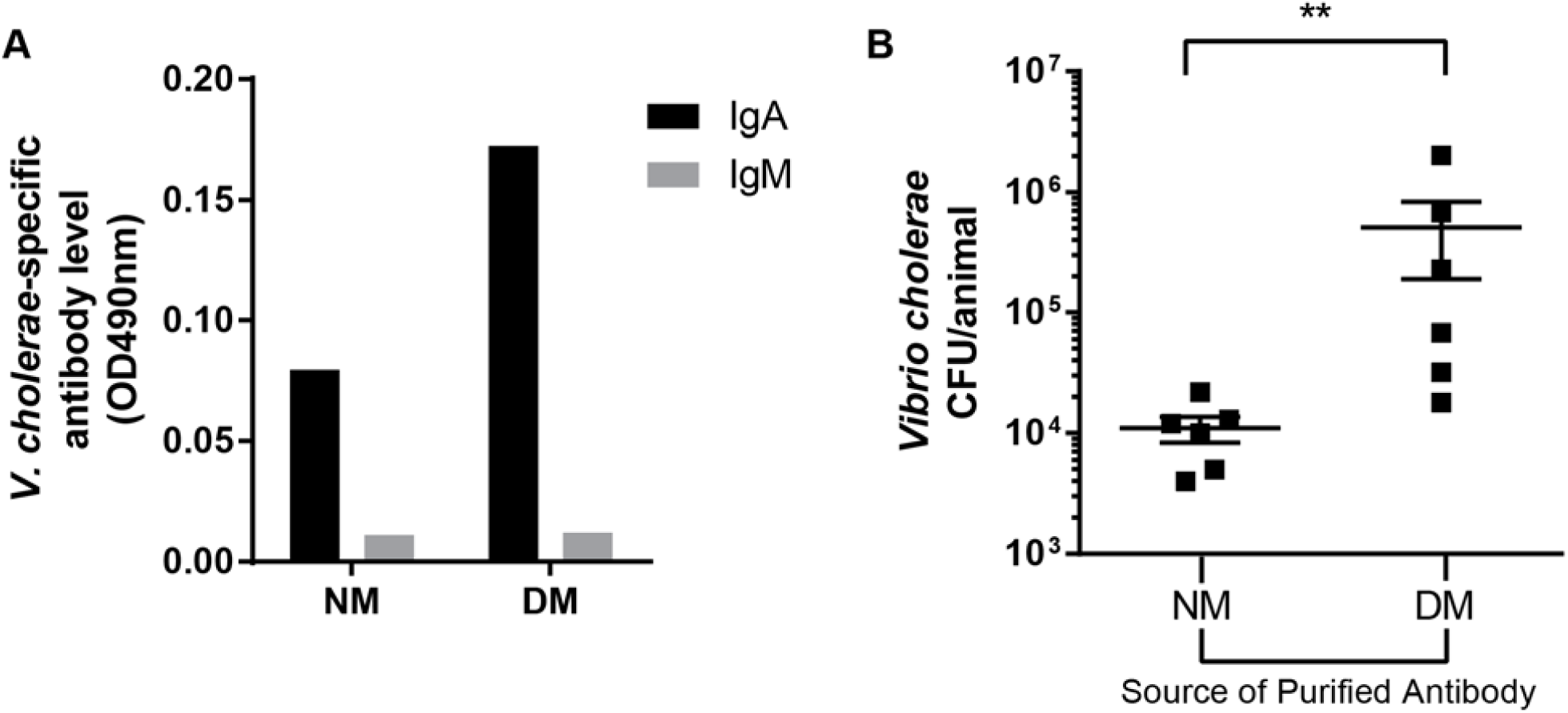
Purified fecal antibody of immunized NM, but not DM animals, can passively protect suckling animals from *V. cholerae* infection. (A) Isotype distribution of pooled, purified fecal antibodies from NM and DM mice. Input was normalized so NM and DM groups received equivalent amounts of IgA. (B) Colonization of suckling CD-1 mice by *V. cholerae* pre-incubated with purified fecal antibody from immunized mice bearing NM and DM microbiomes. **, *P*<0.01, Mann-Whitney *U* test.

To determine whether the NM or DM phenotype would be dominant when the bacterial communities are combined, we immunized mice with *V. cholerae* in the presence of either NM, DM, or NM+DM microbiomes. At 4 weeks post immunization, the NM+DM group showed a low serum vibriocidal titer comparable with the DM group, while the NM group had significantly higher titer levels (Fig. 4A). These data suggested that the dysbiotic microbiome may have a role in suppressing the antibody response against *V. cholerae.*

**FIG 4.**
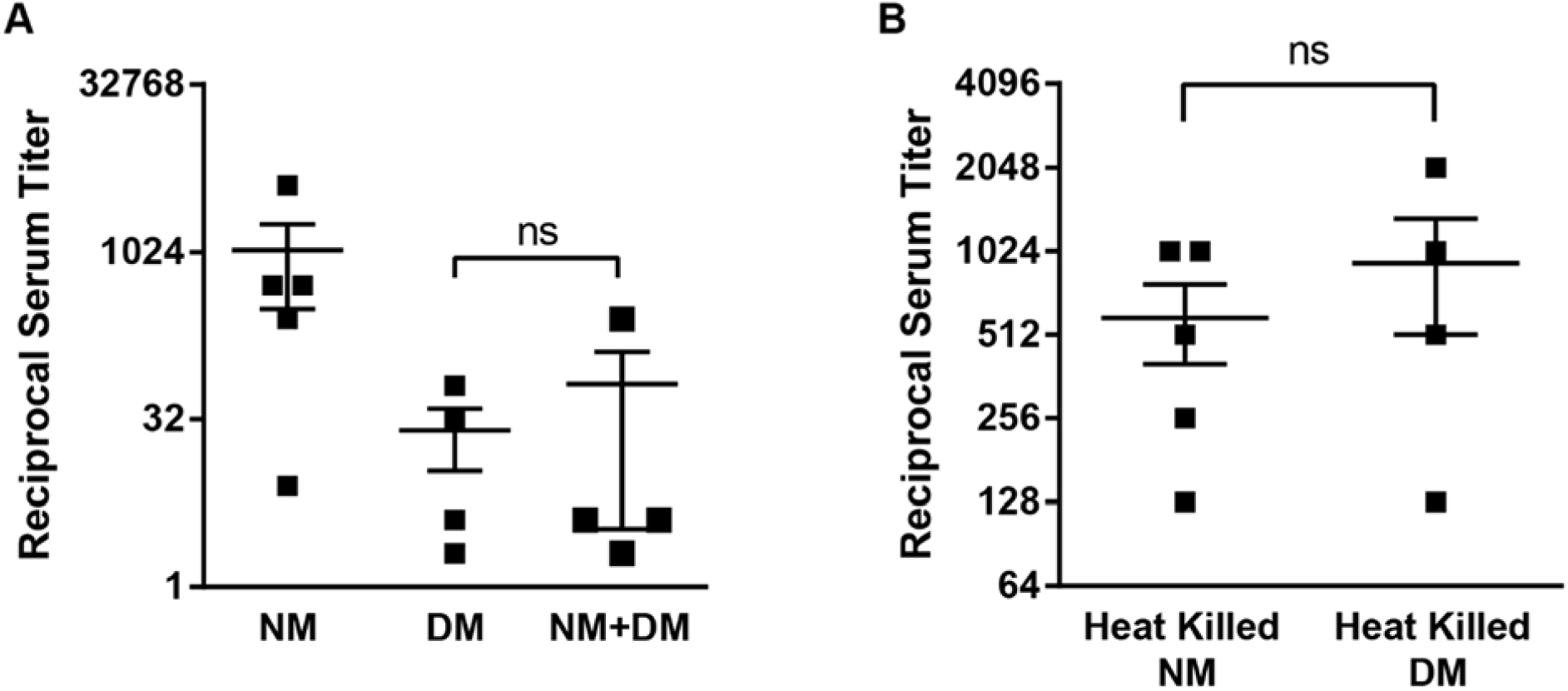
The effect of DM microbes is dominant on immunization outcomes, and requires the presence of live bacteria during immunization. (A) Serum vibriocidal titers 4-weeks post-immunization in CD-1 mice immunized with *V. cholerae* and bearing indicated human model microbiomes. (B) Vibriocidal titers of mice gavaged with indicated heat-killed communities at time of immunization with live *V. cholerae*. ns, *P*>0.05, Mann-Whitney *U* test.

Due to the reduced vibriocidal titer levels observed in the NM+DM group, we wanted to determine whether or not live members of the susceptible community were required to mediate this effect. Accordingly, we heat inactivated all the members of the respective communities and again immunized mice with live *V. cholerae.* When we heat inactivated our bacterial communities, we observed that the serum vibriocidal titer increased in the DM group to similar levels with the NM group (Fig. 4B). These findings suggest that live members of the dysbiotic community are necessary at time of immunization in order to mediate suppression of anti-*Vibrio* antibody protection.

In general, the mechanistic immune responses to OCVs remains to be fully determined. The initial response appears to be driven by TLR-2 interactions that can cause CD4^+^ proliferation, and it has been shown that CD4^+^ T cells are also instrumental in stimulating long-term memory B cell responses (39-42). We therefore further examined the role of CD4^+^ T cells in the context of our DM-mediated suppression of anti-*V. cholerae* antibody responses by depleting CD4^+^ cells in mice co-inoculated with *V. cholerae* and model human microbiomes. Adult CD-1 mice were intraperitoneally injected with anti-CD4 monoclonal antibodies every 4 days during antibiotic treatment. After verifying depletion of CD4^+^ cells by flow cytometry analysis of serum (Fig. 5A-B), animals were gavaged with live defined microbial communities and *V. cholerae* as previously mentioned. Levels of serum anti-*V. cholerae* IgA were severely reduced in both groups (Fig 5C), while there were no significant differences in levels of serum IgG and IgM. Strikingly, vibriocidal titer in the DM group increased to levels comparable to the NM group after CD4^+^ cell depletion (Fig. 5D). This level was also comparable to that observed in NM group without depletion, suggesting that CD4^+^ cells are not required for the development of serum vibriocidal responses. These results suggest that in serum, the primary vibriocidal antibodies are not IgA isotype, and further support for a model whereby the presence of a dysbiotic gut microbiome at time of introduction of antigen leads suppression of subsequent development of specific antibody responses.

**FIG 5.**
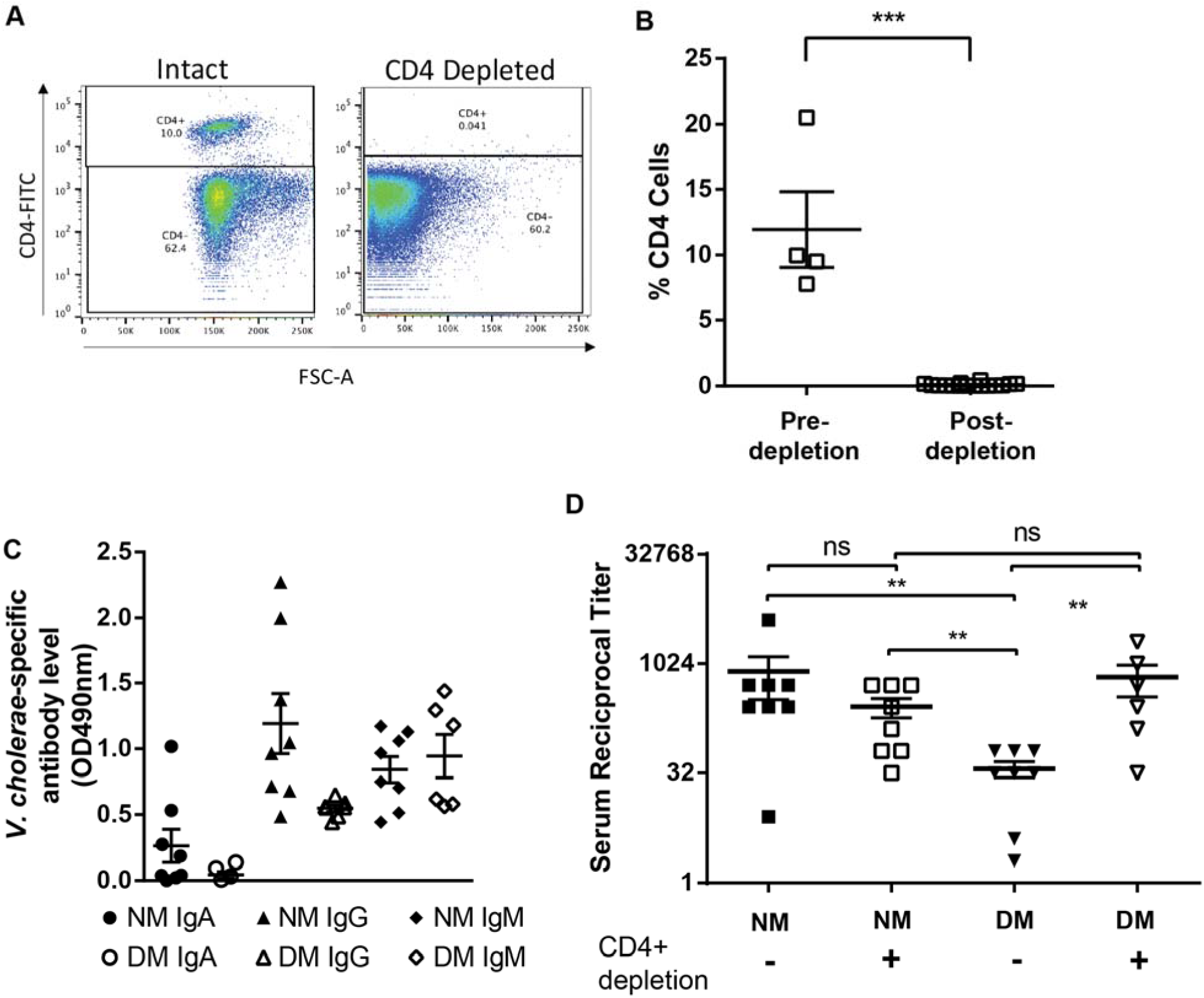
DM community effects are mediated *in vivo* by CD4^+^ cell populations. (A and B) % of CD4 cells pre and 7-days post depletion in blood. (C) Serum antibody levels against *V. cholerae* 4-weeks post-immunization in the presence of indicated human model microbiomes. (D) Comparison of vibriocidal titer levels in NM and DM groups X-weeks after immunization with or without CD4^+^ cell depletion. **, *P*<0.01,***, *P*<0.001, Mann-Whitney *U* test.

## Discussion

Efficacy of vaccination against enteric pathogens has been shown to be highly variable on a geographical and per-study basis, including for rotavirus (43), Salmonella (44), polio(45), and cholera (2). One of the potential reasons for the variability may be due to interpersonal variations in gut microbiomes (46). Previous studies sought to identify the relative abundance of certain species that were either positively or negatively correlated with protection, but were unable to translate these findings into an experimental model of vaccine efficacy. Using microbiome replacement in antibiotic-treated mice, our experiments suggest that the composition of the human gut microbiome at time of immunization suppresses the antibody-mediated immune response against *V. cholerae*. We observed that the presence of the dysbiotic community at time of immunization was sufficient to dampen antibody-mediated protection. Since this required live, as opposed to heat-killed, DM community organisms, these data suggest that certain microbiome structures are actively suppressive of responses to vaccination. There are relatively few studies that have looked at how a community in dysbiosis may alter the humoral immune response. One study shows that *Sutterella* species are capable of degrading the stabilizing peptide of sIgA, leading to decreased levels of IgA (47). The mechanism in our studies is likely different, as our microbial populations are only transiently present, and overall antibody levels are comparable across different model microbiomes.

While in humans, the gut microbiome enters a DM-like state transiently after infectious diarrhea, repeated infection by multiple pathogens, from cholera to pathogenic *Escherichia coli* and rotavirus, means that this state is much more frequently attained in cholera-endemic areas (11, 17, 18). Malnutrition, another common public health concern often co-occurring with recurrent infectious diarrhea, induces a DM-like state for much longer periods, and is refractory to therapeutic nutritional intervention (48). Our DM model community is also similar to the DM-like community state in humans due to its dramatically lower diversity; human microbiomes during fulminant diarrhea and early recovery from diarrhea can be dominated by 99% *Streptococcus* species by relative abundance (17). The definition for what constitutes a truly “healthy” microbiome is not settled. While our NM model community is very broadly reflective of healthy human communities by higher taxonomy levels, interpersonal and temporal intrapersonal variation in healthy individuals can potentially lead to other effects on vaccination outcome not fully captured by our NM-replacement model. More studies with complex human fecal communities will be necessary to probe temporal variations in interactions between the host and the broad range of microbiome structures seen in healthy humans.

Our antibiotic treated adult mouse experimental system is a robust model to study gut microbiota interactions in the host. In contrast to previous studies (22), we are able to transplant actual human microbes into an immune-competent animal system, shortening the loop from initial observations to potentially clinically-relevant conclusions. In contrast, germfree mice have limitations for immune studies; ‘dirty’ mice such as feral and pet store mice display a more mature immune response due to more complete microbial exposure and consequent immune development (49, 50).

To begin teasing apart the host mechanism behind this microbiome-dependent antibody response phenotype, we examined the role of CD4^+^ T cells. Upon depletion of CD4^+^ cells, we observed decreased levels of serum IgA after immunization in both NM and DM mice, potentially indicating decreased seroconversion (Fig. 5C). However, serum vibriocidal titer in DM, CD4^+^-depleted animals increased to levels comparable to the NM mice (Fig. 5D). These data show that CD4^+^ cells are integral in mediating microbiome-dependent changes in an immunization-induced antibody response. These results are surprising as one would expect CD4^+^ T cell depletion to substantially reduce the vibriocidal titer but our data suggests that there are compensatory, non-CD4^+^ mediated mechanisms to aid in seroconversion. A recent clinical study evaluating the efficacy of the oral cholera vaccine Shanchol in human immunodeficiency virus (HIV)-infected individuals demonstrated that while vibriocidal titer was lower in HIV-infected individuals with depleted CD4^+^ T-cell populations there was still seroconversion in 65-74% of the subjects (51). While the study population was not completely depleted of CD4^+^ T cells, it demonstrates vibriocidal titers can be elicited even in a highly-CD4^+^ cell-depleted state, albeit to a lesser degree.

In order to more fully understand the correlations between bacterial communities, *V. cholerae*, and host interactions, more work will need to be done to study the biochemical underpinnings of microbiome-host interaction as it impacts vaccination. The interface between DM microbes and the immune system is yet to be defined; the inability of heat-killed DM communities to influence vaccine outcomes suggest that an active interaction with host tissue, or the production of active compounds *in vivo* are required for this. At the host level, while we investigated the role of CD4^+^ T cells in this phenotype, other immune cell types such as antigen presenting cells may act as more direct intermediaries between host immunity and microbial composition. Helper T cells are integral in stimulating and guiding B cell responses, so it would be beneficial to further define CD4^+^ subsets involved such as follicular helper T cells or regulatory T cells as well as B-cell subtypes.

Taken together, our data advances how bacterial dysbiosis may alter the immune pathways resulting in a weakened humoral response. Ultimately, our studies on the influence of bacterial composition at time of *V. cholerae* immunization will help delineate the contributors to the high variability that occurs in oral cholera vaccines as well as other mucosal vaccines. Our results suggest that the gut microbiome may represent a personalized target for improving vaccination outcome.

## Methods

### Animals

Female CD-1 mice were purchased from Charles River Laboratories, and generally used at 5-9 weeks of age. 4-day old suckling CD-1 mice were purchased from Charles River Laboratories. Animals in the study were treated and housed under specific-pathogen-free conditions. All animal protocols were approved by University of California, Riverside’s Institutional Animal Care and Use Committee.

### Bacterial strains and growth conditions

All human gut commensal strains used are listed in Table S1. Unless otherwise noted, human gut strains were propagated in LYHBHI liquid medium (BHI supplemented to 5g/L yeast extract, 5mg/L hemin, 1mg/mL cellobiose, 1mg/mL maltose and 0.5mg/mL cysteine-HCl). Cultures were then grown in a Coy anaerobic chamber (atmosphere 5% H_2_, 20% CO_2_, balance N_2_) or aerobically at 37°C. All *V. cholerae* strains were derived from the C6706 El Tor pandemic isolate and propagated in LB media with appropriate antibiotics at 37°C.

### Preparation of bacteria for inoculation into antibiotic treated mice

Female adult CD-1 mice were given an antibiotic cocktail *ad libidum* (1 g/L ampicillin, 1 g/L neomycin, and 125 mg/L vancomycin)(52, 53), and for 1 week as described previously with modifications as mice refrained from drinking water with metronidazole (54). 2.5 g/L of Splenda was added as well to make the cocktail more palatable. 3 days prior to gavage with *V. cholerae*, the cocktail was replaced with 2.5 g/L streptomycin and 2.5 g/L Splenda. Each anaerobic human gut bacterium was cultured from glycerol stocks in LYHBHI media for 24 hours at 37°C, and then diluted (1:50) in fresh LYHBHI media. *Enterococcus faecalis* and *Escherichia coli* were grown aerobically in LYHBHI and LB, respectively, for 24 hours at 37°C, and then diluted (1:50) in respective media. After growth for an additional 48 hours, cultures were normalized for density by OD_600_. For inoculation into adult mice, normalized mixtures were prepared so the equivalent total of 300 μL of OD_600_=0.4 culture divided evenly across the respective strains for each community was pooled, centrifuged, and resuspended in LYHBHI. The suspension was prepared so that each mouse received 50 μL of the bacterial community mixture, as well as 50 μL containing ∼5 × 10^9^ *V. cholerae* O1 El Tor C6706. Prior to bacterial introduction, the mice were fasted for 3 hours and then gavaged with 100 μL of 1 M NaHCO_3_, to buffer stomach acid, after which the bacterial communities and *V. cholerae* were inoculated via oral gavage.

### Serum vibriocidal assay

Mouse whole blood was collected via tail vein bleeds using heparinized Caraway collection tubes (Fisher Scientific). Blood was centrifuged at 9,000 × g for 10 minutes, and the serum fraction was isolated and stored at -20°C. The vibriocidal titer measurement was done as previously described with minor modifications (55). In brief, mouse serum was heat inactivated for 30 minutes at 56°C. The heat-inactivated serum was then serially diluted two-fold with phosphate-buffered saline (PBS). Separately, PBS, guinea pig complement serum (Sigma-Aldrich), and ∼ 5 × 10^8^ CFU *V. cholerae* were combined at a ratio of 7:2:1, respectively. The above mixture was then added to the wells containing serially diluted serum and incubated at 37°C for two hours. The resulting dilutions were then plated onto streptomycin (200 μg/mL) LB plates. The vibriocidal titer is the reciprocal of the highest serum dilution which displayed no *V. cholerae* growth.

An additional measure of vibriocidal activity was used by observing the levels of *V. cholerae* remaining after being treated with serum. 25 μL of 5 × 10^7^ CFU *V. cholerae* from an overnight culture was combined with 5 μL serum from the respective groups and 20 μL of PBS. After binding for 1 hour, 5 μL exogenous guinea pig complement was added, and the mixture incubated at 37°C for two hours. Afterward, surviving *V. cholerae* were enumerated by serial dilution and plating on streptomycin LB agar.

### Fecal Pellet Collection

Fresh fecal pellets were collected from mice, weighed, and placed in 600 μL of PBS in a 2.0 mL screw cap tube. The pellets were disrupted by agitation without beads in a bead beater (BioSpec) for 30 seconds at 1400 RPM. 10-fold serial dilutions of the resulting fecal slurry were then plated onto LB agar with streptomycin to enumerate *V. cholerae* colonization.

### Analysis of antibody responses by ELISA

100 μL dense overnight culture of *V. cholerae* grown in LB was plated onto high-binding, clear, flat bottom Costar 96 well plates (Corning, Inc) ELISA plates and allowed to bind overnight. 3% bovine serum albumin (BSA) in PBS was used as a blocking solution. Alternatively, to measure total antibody levels, serum was added at a 1:100 dilution to plates previously coated with unlabeled goat anti-mouse IgA, IgG, IgM (Southern Biotech) and allowed to bind at 37°C for 3 hours. Next, the plates were washed with PBS with 0.001% Tween-20 and PBS. 100 μL of goat anti-mouse HRP conjugated antibodies of either IgA, IgG_1,2A,2B,3_ or IgM (Southern Biotech) were added to 96 well plates at a dilution of 1:4,000 in 3% BSA and incubated overnight at 4°C. After several washes, the plates were developed with the addition of 5 mg o-phenylenediamine dihydrochloride (Thermo Scientific) and stable peroxide substrate buffer (Thermo Scientific); 1 N HCl was used as a stop solution. The plates were read at 490 nm on a Syntergy HTX multi-mode reader (BioTek).

### Passive immune protection assay

Fecal samples from immunized animals bearing the NM and DM communities was collected and processed as described previously. Total IgA/IgM fecal antibody was purified using Protein L magnetic beads according to the manufacturer’s protocol (Pierce Biotech). 50 ng of pure antibody was bound to ∼1.25 × 10^6^ CFU *V. cholerae* and allowed to bind at 37°C for 1 hour. 4-day old suckling CD-1 mice were gavaged with 30-gauge plastic tubing with 50 μL of antibody/*V. cholerae* mixture. After 18 hours of infection, the animals were sacrificed, and intestines homogenized for *V. cholerae* CFU enumeration on selective medium.

### Preparation of heat-killed commensal bacteria

Strains from the NM and DM communities were grown in pure cultures and the bacterial suspension was prepared as previously mentioned. The respective bacterial communities were killed by heating in a heat block for 1 hour at 100°C. Bacterial death was confirmed by plating onto solid media and observing lack of growth.

### *In vivo* depletion of CD4^+^ cells

In order to deplete CD4^+^ cells in vivo, 50 μL of GK1.5 antibody (Bio X Cell) was administered intraperitoneally every four days at a concentration of 100 μg/mL. Depletion of CD4^+^ cells in blood was confirmed using a FACS Canto flow cytometer (BD Biosciences) and FITC rat-anti-mouse CD4 (BD Biosciences). Red blood cell lysis was done using ACK lysis buffer and CD16/32 was used as an Fc block. Analysis was done using Flow Jo (BD Biosciences) and Prism (GraphPad). Mice were treated with ampicillin, neomycin, and vancomycin as previously mentioned. 3 days prior to infection, the mice were placed on streptomycin water alone. The mice were infected with ∼5 × 10^9^ CFU *V. cholerae* and serum vibriospecific ELISAs and vibriocidal assays were performed as previously described.

## Acknowledgments

These studies were funded via a grant from the National Institutes of Health (R35 GM124724). The funders had no role in study design, data collection and interpretation, or the decision to submit the work for publication.

We would like to thank Jonathan Mitchell, Salmasadat Alavi, Jennifer Cho, and Rui Liu for advice and support. We would also like to thank Dr. Meera Nair for helpful discussions as well as Kristina Bergersen and Dr. Emma Wilson for assistance with flow cytometry.

---

**TABLE S1.** Species used.

| Bacterium | Community | Strain Name | NCBI<br>Taxonomy<br>ID |
| --- | --- | --- | --- |
| *Bacteroides vulgatus* | NM | ATCC 8482 | 435590 |
| *Blautia obeum* | NM | ATCC 29174 | 411459 |
| *Clostridium scindens* | NM | ATCC 35704 | 411468 |
| *Enterococcus faecalis* | DM | OG1RF | 474186 |
| *Escherichia coli* | DM | DH5α | 668369 |
| *Streptococcus infantarius* subsp. *infantarius* | DM | ATCC BAA-102 | 471872 |
| *Streptococcus salivarius* subsp. *salivarius* | DM | ATCC 13419 | 1304 |
| *Streptococcus salivarius* subsp. *thermophilus* | DM | DSM 20617 | 1091038 |

